# Evidence for shared conceptual representations for sign and speech

**DOI:** 10.1101/623645

**Authors:** Samuel Evans, Cathy Price, Jörn Diedrichsen, Eva Gutierrez-Sigut, Mairéad MacSweeney

## Abstract

Do different languages evoke different conceptual representations? If so, greatest divergence might be expected between languages that differ most in structure, such as sign and speech. Unlike speech bilinguals, hearing sign-speech bilinguals use languages conveyed in different modalities. We used functional magnetic resonance imaging and representational similarity analysis (RSA) to quantify the similarity of semantic representations elicited by the same concepts presented in spoken British English and British Sign Language in hearing, early sign-speech bilinguals. We found shared representations for semantic categories in left posterior middle and inferior temporal cortex. Despite shared category representations, the same spoken words and signs did not elicit similar neural patterns. Thus, contrary to previous univariate activation-based analyses of speech and sign perception, we show that semantic representations evoked by speech and sign are only partially shared. This demonstrates the unique perspective that sign languages and RSA provide in understanding how language influences conceptual representation.

## Introduction

Conceptual knowledge is fundamental to human cognition. Recent evidence suggests that conceptual representations are flexible and contextually defined^1,2^. Does the language that we use influence the nature of stored conceptual representations? If this is the case, we might predict that languages that differ most in structure, such as sign and speech, would show the greatest divergence between conceptual representations. Sign languages are visuo-spatial natural languages that are distinct from surrounding spoken languages. Hearing people with signing deaf parents are bilingual in sign and speech. These individuals offer a unique insight into the influence of both modality and bilingualism on semantic processing.

Semantic cognition engages a distributed left lateralised fronto-temporo-parietal network^3,4^. Strong evidence for modality independent neural representations comes from studies using multivariate cross-classification of functional Magnetic Resonance Imaging (fMRI) data that show that neural patterns elicited by an item in one modality (e.g., pictures) can predict patterns for the same item presented in a different modality (e.g., spoken words). These studies have identified common patterns within hearing participants for pictures, identifiable sounds and spoken and written words in the inferior temporal, parietal and prefrontal cortex^5–7^. Data from patients with semantic dementia also suggest an important role for the inferior anterior temporal lobe in semantic cognition, as a modality independent “hub”^2^. However, studies of the influence of modality on semantic processing in hearing participants might reflect the eliciting of common oral language representations via visual and auditory stimuli ^8,9^. Therefore, contrasting representations evoked by sign and speech in hearing sign-speech bilinguals, offers a stronger test of the influence of modality on semantic processing, whilst also providing a unique perspective on bilingualism.

How multiple languages are represented in a single brain is still not clear. Evidence for shared representations comes from cross-linguistic priming^10^ and stroop-type tasks^11^ in spoken language bilinguals. However, evidence from word association and translation tasks suggest different or only partially overlapping semantic representations between languages^12,13^. At the neural level, fMRI studies show both common and language specific activity elicited by the different languages of bilinguals^14–18^. In these studies, the relative contribution of phonology, semantics and syntactic processing has not been explicitly differentiated. Studies of bilinguals to date have typically investigated across language representations ***within-modality***, e.g. from speech to speech, or text to text. Only one study has attempted the stronger test of cross classifying between *both* language *and* modality. They found it was not possible to cross-classify neural patterns for individual written and heard words across different spoken languages^19^.

Sign and speech are conveyed in different modalities. Despite this, univariate analyses of speech and sign perception reveal substantially overlapping brain networks^20–26^. However, to date, the similarity of neural patterns evoked by individual signs and spoken words has not been quantified. Here, using representational similarity analyses^27^, we assess the evidence for shared and language specific representations of individual conceptual items and semantic categories, for speech and sign in hearing, early sign-speech bilinguals. Our findings provide evidence for shared semantic representations at the level of categories, but not for individual conceptual items. This suggests that visuo-spatial languages and spoken languages evoke subtly different conceptual representations.

## RESULTS

In the scanner, hearing early sign-speech bilinguals were presented with 9 conceptual items from the 3 semantic categories: fruit, animals or transport. Each item was presented as a sign or as a spoken word and was produced by a male or a female language model (Fig. 1a). Participants were instructed to press a button to detect occasional items, 8% of the trials, that were not from one of the 3 target categories (Fig. 1b). Performance in the scanner indicated that participants were fully engaged with the semantic monitoring task (see Supplementary Information 1). A univariate GLM analysis indicated that speech and sign language engaged similar fronto-temporal networks, consistent with previous studies^20–24^ (see Supplementary Information 2).

**Fig. 1.**
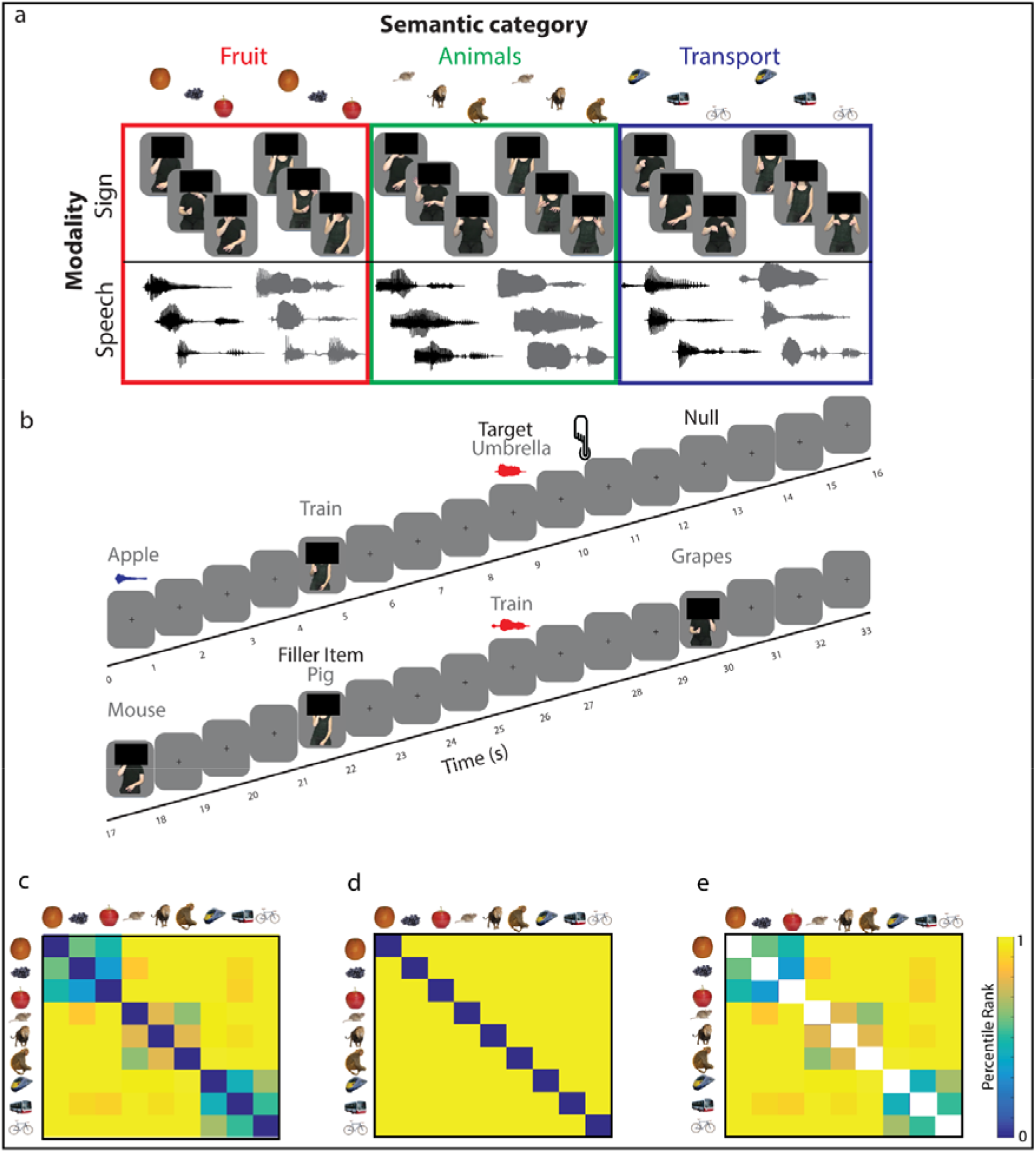
Stimuli, experimental design and semantic models. (Fig. 1a) Early sign-speech bilinguals were presented with 9 conceptual items that belonged to 3 semantic categories: fruit, animals and transport. Items were presented as signs and spoken words and were produced by male and female language models. Video stills and oscillograms are shown for the signs and spoken words respectively. Please note that the faces of the language models have been obscured to comply with the policy of BioRxiv. Participants saw the faces of the signers. (Fig. 1b) Within the scanner, participants attended to speech and sign and pressed a button to identify items that were not in one of the three target categories (e.g., umbrella). The dissimilarity between neural patterns evoked by the signs and spoken words were tau-a correlated with different theoretical models. These models included (Fig. 1c) a semantic feature model derived from the CSLB concept property norms^28^. The color bar reflects the degree of semantic dissimilarity between items. This semantic feature model can be decomposed into two independent components: (Fig. 1d) An item-based dissimilarity model that predicts that each item is uniquely represented, e.g., an ‘apple’ is more dissimilar to other items than to itself and does not predict any broader semantic relatedness between items and (Fig 1e) a category-based model in which the between-item similarities are predicted by the semantic feature model, but where the within-item similarities are not tested. White squares in this model indicate comparisons that were excluded.

### Shared semantic representations for speech and sign

Our criteria for identifying shared semantic representations for speech and sign were as follows. First, using a searchlight analysis, we identified regions in which there were reliably positive distances (see methods) between items ***within-modality*** (e.g. averaging the speech-speech distances and the sign-sign distances). We calculated distances only between items from the different language models (e.g. different speakers and signers respectively) to exclude similarities driven by low-level perceptual properties. In the identified regions, we then tested for ***shared semantic representations*** applying the following criteria: (A) a significant fit to the semantic feature model in the ***within-modality*** distances (e.g. both the across speaker, speech-speech, and the across signer, sign-sign, distances) and (B) a significant fit of the semantic feature model to the ***across-modality*** distances (e.g. speech-sign and sign-speech distances). We also expected, (C) no evidence of a difference in strength of fit to the semantic model between speech and sign, (D) no fit to a model predicting greater distances between items from a different, as compared to the same speaker, in the speech-speech distances, or from a different, as compared to the same signer, in the sign-sign distances and (E) no fit to a model predicting sensitivity to the iconicity of sign, a perceptual feature present in sign but not speech.

Reliable within-modality distances were identified in six clusters (Fig. 2a): **(1)** in bilateral V1-V3 and the LOC [−14 −96 10], **(2)** the right anterior superior temporal gyrus [58 −4 −2], **(3)** the left anterior superior and middle temporal gyrus [−60 −10 −2], **(4)** the right middle temporal gyrus and MT/V5 [52 −68 6], **(5)** the right insular [36 −12 14] and **(6)** the left posterior middle and inferior temporal gyrus (left pMTG/ITG) [−48 −62 −6].

**Fig. 2.**
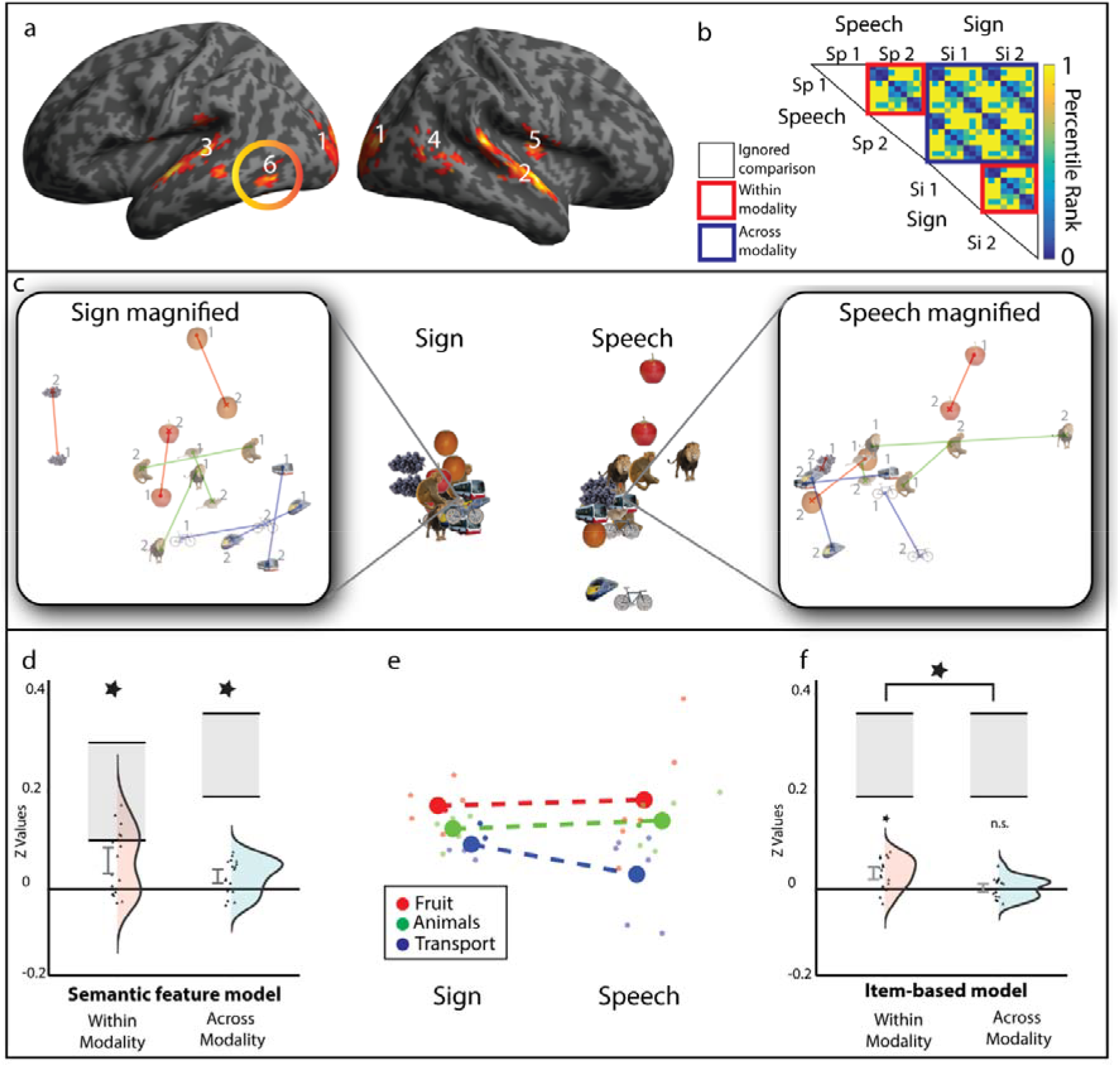
Shared semantic representations for speech and sign. (Fig. 2a) A searchlight analysis identified brain regions containing positive *within-modality* representational distances, thresholded at p < 0.005 peak level, FDR corrected at q < 0.05 at the cluster level. These regions are numbered according to the text in the results section. (Fig. 2b) Representational distances in these regions were Tau-a correlated with the semantic feature model within- and across-modality. The red boxes illustrate the within-modality distances, with the upper red box testing for abstracted speech representations (e.g. from speaker 1 to 2), and the lower red box testing for abstracted representations for sign (e.g. from signer 1 to 2). The blue box contains all across-language distances. Each 9×9 submatrix of dissimilarities is predicted from the semantic feature model (Fig. 1c). White boxes are comparisons excluded from the analysis. The color bar reflects the predicted strength of dissimilarity. Plots (Figs. 2c-f) show the response in cluster 6, the left pMTG/ITG. (Fig. 2c) shows the non-metric MDS representation of the response in left pMTG/ITG: the left panel shows within sign distances magnified to make the representational structure clearer and the right panel shows the equivalent speech representations. In these magnified images, lines connect the same conceptual item produced by each speaker or signer, marked as speaker/signer 1 or speaker/signer 2 on the figure. (Fig. 2d) In the left pMTG/ITG, there was a significant fit to the semantic feature model in both the within- and across-modality distances. Violin plots show distributions and individual data points for the z transformed values, including the 90% confidence interval and the noise ceiling (grey rectangle). The relative contribution of item-based (Fig. 1d) and category-based (Fig. 1e) to this fit was assessed. This showed there to be a significant fit to the category-based model both within- and across-modality, without evidence of a difference in fit when they were compared with one another. The MDS representation (Fig. 2e) showing the mean centroid of each category within each modality for fruit (red), animals (green), blue (transport), with dashed line connecting centroids across-modality, highlights the within and across-modality category-based dissimilarity. Plot (Fig. 2f) demonstrates that the item-based model was a significant fit to the within-modality, but not across-modality distances, and that the item-based model was a better fit to the within- as compared to across-modality distances.

Only the response in the left posterior middle and inferior temporal gyri (pMTG/ITG) cluster was consistent with shared semantic representations (see Fig. 2a cluster 6; Supplementary Information 3 for full details). In this cluster, there was a significant fit to the (A) ***within-modality*** semantic feature model (t (16) = 3.622, p = 0.001, d_z_ = 0.879, Fig 2d) and (B**) *across-modality*** semantic feature model (t (16) = 3.076, p = 0.004, d_z_ = 0.746, Fig 2d). Whilst there was (C) no evidence for differential sensitivity in the encoding of semantics for speech and sign (t (16) = 0.400, p = 0.694, d_z_ = 0.097), (D) no sensitivity to the acoustic or visual features associated with speaker (see model in Fig. 3e) or signer identity (see model in Fig. 4e), both ps > 0.063, or (E) no influence of the iconicity structure of sign in the sign-sign or across-modality distances, both ps > 0.106 (see Supplementary Information 4 and Supplementary Fig. 2).

**Fig. 3.**
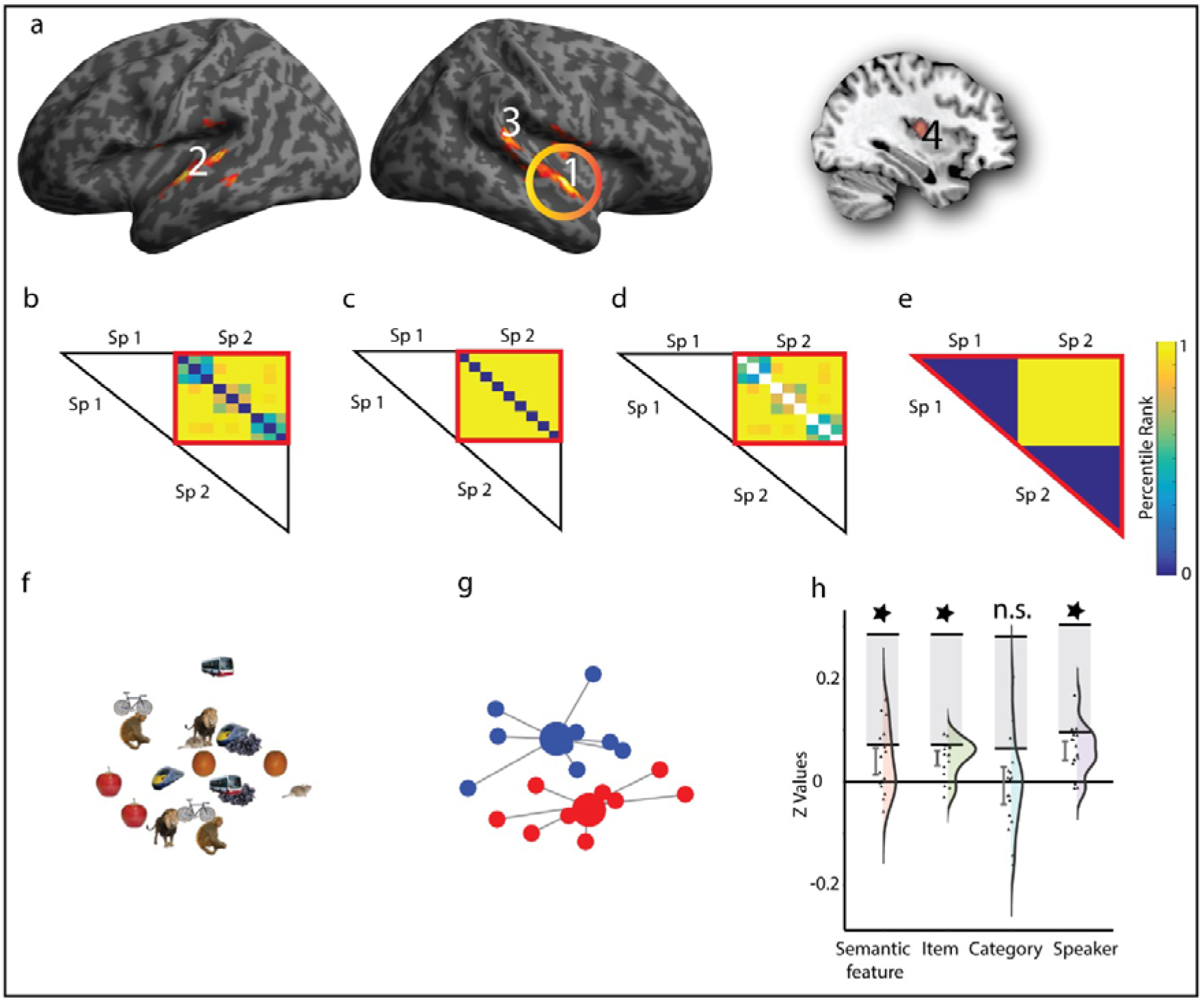
Speech-specific neural responses. (Fig. 3a) A searchlight analysis identified regions with greater representational distances for speech compared to sign, thresholded at p < 0.005 peak level, FDR corrected at q < 0.05 at the cluster level. Clusters are numbered according to the text in the results section. Models (Figs. 3b-e) show the within speech models that were tested: (Fig. 3b) Within-speech semantic feature model, (Fig. 3c) Within-speech item-based model, (Fig 3d) Within-speech category-based model and (Fig. 3e) Between-speaker model. All models (Figs. 3b-d) test dissimilarities across speaker (e.g. from speaker 1 to 2) in order to identify representations abstracted from perceptual features. Color bar reflects predicted strength of dissimilarity. White boxes are comparisons excluded from analysis. Plots (Figs. 3f-h) show the response in cluster 1, the right anterior STG: (Fig. 3f) Shows the non-metric MDS solution and (Fig. 3g) the same solution highlighting speaker identity encoding. Large circles represent the centroids for items from speaker 1 (red) and speaker 2 (blue). Smaller circles represent the observed response for each item. Grey lines connect each item to centroid. (Fig. 3h) Violin plots show model fits for z transformed values for each model, with distributions and individual data points and 90% confidence intervals and noise ceiling (grey box shown). This shows a significant fit to the semantic feature model, driven by item-based rather than category-based similarity structure and additional sensitivity to speaker identity, consistent with abstract spoken word form representations rather than modality specific semantic processing.

**Fig. 4.**
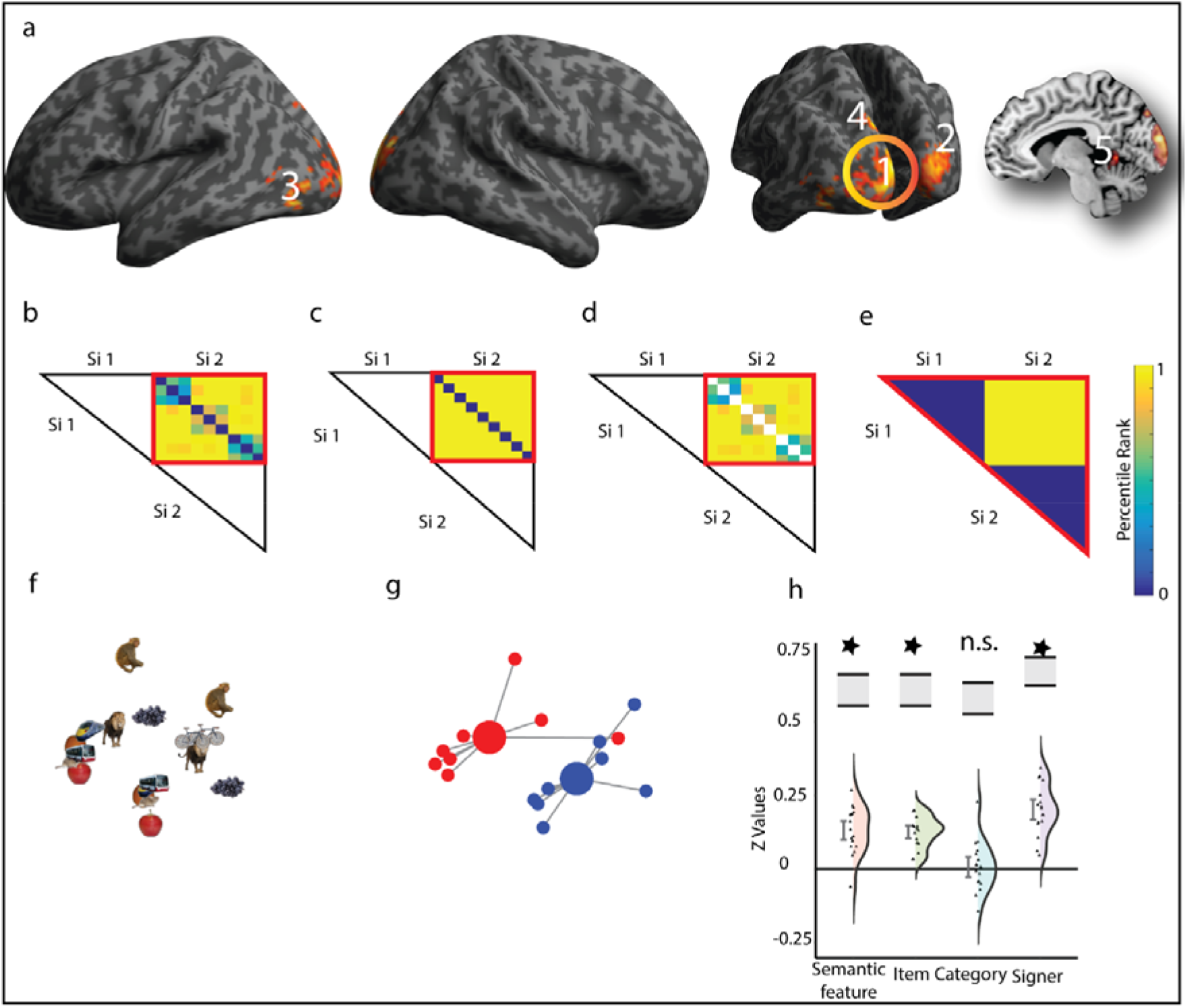
Sign specific neural responses. (Fig. 4a) A searchlight analysis identified regions with greater representational distances for sign compared to speech, thresholded at p < 0.005 peak level, FDR corrected at q < 0.05 at the cluster level. Clusters are numbered according to the text in the results section. Models (Figs. 4b-d) show the within sign models: (Fig. 4b) Within-sign semantic feature model, (Fig. 4c) Within-sign item-based model, (Fig. 4d) Within-sign category-based model and (Fig. 4e) Between-signer model. Models (Figs. 4b-d) test dissimilarities across signer (e.g. from signer 1 to 2) to identify representations abstracted from perceptual features. Color bar reflects predicted strength of dissimilarity. White boxes are comparisons excluded from analysis. Plots (Figs. 4f-h) show responses in cluster 1, the left V1-V3. (Fig. 4f) Shows the non-metric MDS solution and (Fig. 4g) the same solution highlighting the signer identity encoding in the left V1-V3 cluster. Large circles represent the centroids for items from signer 1 (red) and signer 2 (blue). Smaller circles represent the observed response for each item. Grey lines connect each item to centroid. (Fig. 4h) Violin plots show model fits for z transformed values for each model fit, with distributions and individual data points and 90% confidence intervals and noise ceiling (grey box shown). Plots show a significant fit to the semantic feature model, driven by item-based rather than category-based similarity structure and an additional sensitivity to signer identity within the left V1-V3, consistent with abstract sign form representations rather than modality specific semantic processing.

The fit of the semantic feature model (Fig. 1c) can be further decomposed into item-based dissimilarity (Fig. 1d) and category-based dissimilarity (Fig. 1e). For ***within-modality*** distances, the left pMTG/ITG region showed a significant fit to both the semantic category (t (16) = 1.980, p = 0.033, d_z_ = 0.480) and item-based model (t(16) = 4.185, p = 3.50 × 10^−4^, d_z_ = 1.015). The critical analyses ***across-modality***, indicated that the category-based model showed a significant fit to the data (t (16) = 2.509, p = 0.012, d_z_ = 0.608), whereas the item-based model did not (t (16) = 0.475, p = 0.321, d_z_ = 0.115). There was no evidence of a difference in the strength of fit to the category model in the ***within-modality*** as compared to the ***across-modality*** distances (t (16) = 0.135, p = 0.894, dz = 0.033), suggesting that semantic categories were represented equally robustly within- and across-modality. By contrast, the item model was a significantly better fit to the ***within-modality*** than the ***across-modality*** distances (t (16) = 3.376, p = 0.004, dz = 0.819, Fig. 2f), providing strong evidence that item-based representations are less robustly encoded across-modality.

Together, these results suggest that semantic category structure drives the commonality between activation patterns for sign and speech in left pMTG/ITG. Indeed, this can be seen in the Multidimensional Scaling (MDS) solution (Fig. 2c) used to visualise the similarity structure of the Representational Dissimilarity Matrix (RDM). Fig. 2e illustrates the similar ordering of the category centroids both within and across each modality.

### Modality specific representations

Using a searchlight analysis, we tested for regions in which the average of the speech-speech distances were greater than the sign-sign distances and vice versa. This identified speech-specific and sign-specific processing regions. Within these regions we tested for ***modality specific semantic representations*** evidenced by (A) a fit to the semantic feature model (Fig. 1c) and (B) a fit to the semantic category model (Fig. 1e) in the speech-speech or sign-sign distances for speech or sign respectively and (C) no evidence of a fit to the speaker or signer identity model (see the models in Fig. 3e and 4e).

#### Speech specific responses

For speech, the searchlight analysis revealed four clusters: **(1)** right anterior STG extending to the temporal pole [58 −4 −2], **(2)** left anterior STG [−56 −8 2], **(3)** right posterior STG/STS [58 −34 18] and **(4)** right putamen and insula [30 −10 10] (see Fig. 3a). Within these regions, we tested for speech specific semantic representations adjusting the critical alpha level to p < 0.013 to account for tests in four clusters. In one of the four clusters, the right anterior STG [58 −4 −2] (Fig. 3a, cluster 1), there was a significant fit to the semantic feature model (t (16) = 2.529, p = 0.011, d_z_ = 0.613, see Fig 3b and Fig. 3h). This was driven by a fit to the item-level model (t (16) = 5.229, p = 4.14 × 10^−5^, d_z_ = 1.268, see Fig. 3c and Fig. 3h). This region was additionally sensitive to the acoustic differences between speakers (t (16) = 5.330, p = 3.39 × 10^−5^, d_z_ = 1.293, see Fig. 3e and Fig. 3h) suggesting the presence of speech form representations rather than speech selective semantic representations (see Fig. 3f and Fig. 3g for MDS solution highlighting speaker-based similarity). None of the four regions showed a response consistent with speech specific semantic representations, as the category-based model (Fig. 3d) was not a significant fit in any region (all ps > 0.110, see fit to the speaker model in the right STG in Fig. 3h).

#### Sign specific responses

Greater representational distances for sign than speech were identified in five regions: **(1)** a cluster spreading across left V1-V3 [−6 −98 16], **(2)** a cluster within right V1-V3 [22 −90 16], **(3)** a cluster in the left LOC and MT/V5 [-44 −80 −6], **(4)** left superior occipital gyrus and superior parietal lobule [−10 −84 42] and **(5)** left lingual gyrus spreading to the cerebellum [−4 −48 −8] (see Fig. 4a). Within these regions, we tested for sign-specific semantic representations, adjusting the critical alpha level to p < 0.010 to account for tests in five clusters. Analogous to the findings for speech, the response in these regions was not consistent with sign-specific semantic representations, as the category-based model was not a significant fit in any region (all ps > 0.037). The response in clusters in the left V1-V3 and right V1-V3 cluster were consistent with sign form representations characterised by a significant fit to the semantic feature model (both ps < 3.10 × 10^−5^) but driven by item-based encoding (ps < 1.34 × 10^−7^) with additional sensitivity to signer identity (both ps < 3.29 × 10^−7^, see Fig. 4).

## DISCUSSION

On the basis of univariate analyses of fMRI data it has been assumed that the same underlying semantic representations support the perception of spoken and signed languages ^29^. We tested this assumption, using RSA, to quantify the similarity of neural patterns evoked by the same conceptual items presented as BSL and spoken British English: two languages that differ in their modality of expression. We tested for similarity at the level of individual items and semantic categories. Shared category representations, that were abstracted from surface acoustic and visual form, were found in the left pMTG/ITG. In this region, both individual items and categories were encoded within-modality. Across-modality, we found evidence for common coding of semantic categories. We did not detect evidence of common item-level representations across modalities. Furthermore, item-level encoding was significantly stronger within- as compared to across-modality. In sign-specific and speech-specific areas, mainly in visual and auditory primary and association cortices respectively, there was evidence for modality specific item-based representations. In these regions, we did not see evidence for category-based structure and the representations retained sensitivity to auditory and visual features, suggestive of phonological word and sign form representations rather than language specific semantic representations. Taken together, our data are consistent with shared semantic representations between speech and sign, at only a broad level of semantic specificity. In the following sections, we discuss the implications of these findings.

### Shared semantic representations in pMTG/ITG

We identified shared representations for semantic categories in sign and speech within the left pMTG/ITG. This is consistent with studies showing common category representations for the same items presented as pictures, environmental sounds, and spoken and written words in this region ^5,7^. Indeed, activation of the left pMTG/ITG is associated with the extraction of meaning from both the auditory and visual modalities. For example, it is activated when reading words^30^, in the perception of semantically ambiguous speech^31^ and during sign language perception ^25,26,32^.

Common semantic coding for sign and speech was limited to category representations and there was no evidence for direct correspondences between individual spoken words and signs. Partially shared semantic representation between languages is consistent with computational models of bilingualism, such as the Distributed Feature Model^33^. These models predict a single semantic store, in which each language weights semantic features independently^13,33,34^. The factors contributing to differing weights between signed and spoken languages may be greater than, and different to, those contributing to divergence between spoken languages. Studies of spoken language processing show that lexical-semantic access is affected by the phonological structure of the lexicon. For example, words from dense phonological neighbourhoods activate semantic representations less strongly^35^ due to cascading activation between phonology and semantics^36^. Indeed, many computational models of speech processing do not make distinctions between form and meaning^37^. Similar architectures have been suggested for sign processing^38^. As natural languages, signed and spoken languages have very different phonologies and phonological neighbourhoods. This might affect the strength and structure of semantic activation within sign and speech lexicons, with the possible result of reducing the commonality of conceptual representations between the languages.

Another possibility is that the influence of greater iconicity found in sign languages^39^ may reduce the degree of similarity between semantic representations of sign and speech. However, this is an unlikely explanation for the lack of item-level correspondences between individual words and signs in the current dataset, as we did not observe an effect of iconicity in the response in the left pMTG/ITG. There are, however, more opaque form-meaning links that differ across speech and sign. For example, the handshape “I” (extension of the little finger alone) denotes a number of BSL signs that have negative connotations: bad, wrong, awful, poison^40^. Similarly, English words beginning with “gl” are often associated with light of low intensity: gleam, glow, glint, glimmer, glint^39^. Canonical signs can also carry additional layers of meaning that allow communication of the size, location, movement and other features of the referent; aspects of meaning that cannot be communicated by the paralinguistic features of the voice. Again, these features may fundamentally change the nature of semantic representation. These potential explanations for the lack of item-level correspondences need to be tested in future. For example, based on these findings, we might predict differences in the representation of specific semantic categories, for example, representations for tools might be expected to differ between unimodal (e.g. speech-speech) and bimodal (e.g. sign-speech) bilinguals, on the basis that signs evoke greater specificity in the semantic features associated with how they are handled.

An alternative explanation is that the absence of shared item-level correspondences reflects the finer spatial scale of neural representations for individual items which might be beyond the resolution of fMRI^41^. However, this would seem unlikely given the identification of within-modality item-level encoding. Equally, it might also reflect our methodological choices. We asked participants to monitor for category rather than item-level distinctions^42^. We decided to use a category-based task to maximise the likelihood of finding commonality between the languages, which we assumed would be more robust at a broader level of semantic specificity. Another possibility is that we did not have a high enough signal to noise ratio in areas in which across-modality item level representations might be expected. A posterior-anterior gradient of function has been suggested within the inferior temporal cortex that reflects a wider-to-narrower window of semantic specificity^2,43^. The anterior inferior portion of the inferior temporal cortex is particularly susceptible to signal drop out. Hence, the absence of shared item-level encoding might reflect reduced signal quality in this region. However, tSNR maps for our data indicate relatively good signal quality in most of the anterior inferior temporal cortex (see Supplementary Information 3). Furthermore, drop out in the anterior inferior ATL was similar to that found in the left pMTG/ITG and the superior ATL, regions in which we found significant representational structure. We chose not to use a dual echo sequence to mitigate against drop out ^44^, as our sequence was optimised for signal quality in the posterior temporal cortex, the region most consistently activated by both sign and speech in previous univariate studies. Future studies using dual echo sequences and item-level discriminative tasks are necessary to exclude the possibility that these methodological details obscured identification of item-level correspondences in this study.

### Modality specific representations

Greater representational structure for speech, than sign, was found in the bilateral superior temporal cortex and the right insula. Within these regions, only a cluster in the right anterior superior temporal cortex was a significant fit to the semantic model. This was shown to be driven by the encoding of individual spoken words. A role for the anterior superior temporal cortex in representing the identity of spoken words is consistent with studies in which the intelligibility of speech has been parametrically varied or contrasted with non-speech sounds^45,46^ and the suggestion that spoken word representations are detected in the more superior portion of the ATL ^2^. This region was additionally sensitive to speaker identity, suggesting that spoken word forms and speaker characteristics are jointly encoded. This is consistent with a role for the right anterior superior temporal cortex in representing speaker identity^47^ and weak joint sensitivity to spoken word and speaker identity in the right superior temporal cortex^48^. The fact that representations of spoken word forms were identified in the right, but not left anterior STG, is unexpected. One possibility is that it is due to the greater involvement of right hemisphere structures in language processing in early bilinguals^49^.

Regions containing greater representational structure for sign, than speech, were found in the bilateral occipital cortices, as well as in the left superior parietal lobule. This is consistent with the greater visual and body-space processing demands of sign language perception ^29^ and the growing evidence for superior parietal cortex involvement in sign perception and production^50^. As for speech, a subset of regions showing greater representational structure for sign than speech showed a significant fit with the semantic model, and this was driven by item-level encoding, consistent with visual sign form representations. Paralleling the findings for speech, a number of these regions also exhibited a joint sensitivity to the identity of the sign and the signer.

### Conclusions

For the first time, we quantified the similarity of neural representations for the same conceptual items presented as sign and speech. We found similarity between conceptual representations, at the category level, in the left pMTG/ITG. We did not find evidence for regions in which there were direct one-to-one mappings between individual spoken words and signs. This may suggest that sign and speech share partially, but not fully, overlapping semantic representations. This result is unexpected. Evidence to date has led researchers, including ourselves, to propose extensive similarity in the neural processes underlying sign and speech ^29^. Our findings suggest the need to rethink this assumption and highlight the unique perspective that sign language can provide on language processing and semantic representation more broadly.

## Supporting information

All supplementary materials

## Online Methods

### Participants

Ethical approval was granted by the UCL ethics committee. Data were collected from 18 right handed early sign-speech bilinguals with no known neurological, hearing or language learning impairments. One participant’s data was removed from the set due to an incidental finding, leaving a final data set of 17 participants (Mean age=33; range 20-52 years; female=12). Fifteen participants learned British Sign Language (BSL) from a deaf parent and two from an older deaf sibling. Two of the participants who learned sign language from a deaf parent did not learn BSL from birth; one, learned AUSLAN from birth and learned BSL from the age of twenty-one, the other, was exposed to another sign language from birth, before learning BSL from 3 years of age. As a group the participants self-reported excellent signing ability (mean = 6/7, SD= 0.86, range = 4-7).

### Stimuli

Stimuli consisted of nine core items for which neural responses were analysed. Each core item was presented 48 times across the whole experiment, in different modalities (sign/ speech) and by different models (male/ female) (see ‘paradigm’ for more details). These nine items belonged to three categories: fruit (orange, grapes and apple), animals (mouse, lion and monkey) and transport (train, bus and bicycle). Items within each category were similar and were distinct from other categories on the basis of their semantic features, as evidenced by the CSLB concept property norms^28^ (see Fig. 1c). Items were chosen to ensure that the categories were matched for age of acquisition (fruit M = 3.78; animals M = 4.52; transport = 4.04), imageability (fruit M = 618; animals M = 610; transport M = 640), familiarity (fruit M = 566; animals M = 521; transport M = 551) and the number of syllables and phonemes in spoken English^51–54^. In addition, we ensured that the BSL equivalents of the spoken words were matched across category for handshape, location, movement and handedness, and that iconicity^55^ was similar across categories (fruit M = 3.80; animals M = 3.92; transport M = 4.23; 1 low - 7 high iconicity).

Speech samples were recorded by a male and female Southern British English (SBE) speaker in an acoustically shielded booth with 16-bit quantisation and a sampling rate of 22050 Hz using Adobe Audition. Spoken words were excised at the zero crossing point. They were then filtered to account for the frequency response of the Sensimetric headphones used in the scanner (http://www.sens.com/products/model-s14/) and the overall amplitude was Root Mean Square (RMS) equalised to ensure a similar perceived loudness (see Fig. 1a for oscillograms). The mean duration of the auditory stimuli for the core items was 558ms (range = 323-865 ms), these sounds were similar in duration across semantic categories (fruit M = 573 ms; animals M = 575 ms; transport M = 533 ms) and gender of the speaker (male M = 557 ms; female M = 564 ms). The phonetic distance between each of the spoken words was calculated using the Levenshtein distance^56^. This was achieved by calculating the number of phoneme insertions, deletions and/or substitutions necessary to turn one word into the other, divided by the number of phonemes in the longest word. The absolute value of the difference in Levenshtein distance between each item was calculated. These distances did not correlate with the semantic feature distances (r = 0.063, n = 36, p = 0.713), hence semantic structure was not confounded with phonetic structure.

The BSL signs were all common variants in southern England as shown in the BSL SignBank^57^ (http://bslsignbank.ucl.ac.uk/dictionary/). Signs were recorded with a Sony Handycam HDR-CX130 on a blue background by a male and a female deaf native signer with a sampling rate of 50 fps and an aspect ratio of 1920×1080. The blue background was keyed out and replaced with a dark grey background. Videos were down-sampled to 30 frames per second and a resolution of 960 × 540 with Adobe Premiere for presentation in the scanner. All signs were produced with corresponding BSL mouthing. The signs were recorded in isolation such that the hands returned to a neutral position resting on the knees between each sign. During editing, the start and end-points of a sign were identified as a ‘hold’ (very brief pause in movement of the hands) to remove the transitional movement into and out of the neutral hands on the lap. Still frames of the hold points at the beginning and end of each sign, with duration of 333ms, were inserted to ensure that the signs were easily perceived in the scanner. The mean duration of the sign stimuli was 1107ms (range = 867-1400ms). The signs were similar in duration as a function of semantic category (fruit M = 1079ms; animals M = 1055ms; transport M = 1128ms) and gender of the signer (male M = 1087ms; female M = 1086ms).

An iconicity dissimilarity measure^55^ for the signs was calculated by taking the absolute value of the difference between ratings of each item with every other. These distances did not correlate with semantic feature similarity (r = −0.126, n =36, p=0.465), hence semantic structure was not confounded with iconicity.

Participants were shown 36 additional items in the scanner to facilitate a semantic monitoring task (see Fig. 1b) for which neural activity was not analysed. The additional items consisted of 18 items from outside the categories of fruit, animal and transport, e.g. buildings, clothes, furniture and tools, which were included as target filler trials. Plus, an additional 18 non-target filler trials, 6 per category, of other types of fruit, animals or transport that were included to reduce habituation to the nine core items (see ‘Paradigm’ below for details of number of presentations). Each individual filler item was produced by only one of the speakers or signers, with the number of items from each speaker and signer balanced. The full set of stimuli are available here: https://osf.io/ek8ty/.

Prior to scanning, participants were familiarised with the signs and spoken words. Participants saw each sign stimulus and heard each word produced by both sign and speech models and were required to name each item in spoken English. They were shown all core items, target and non-target fillers. Sign recognition was high (core items: mean = 17/18, min = 15/18, max = 18/18; filler items: mean = 32/36, min = 21/36, max = 35/36). On the very few occasions that participants interpreted a sign as a non-intended English word, due to regional variations in signs, participants were told the intended spoken label and asked to repeat it. They were then retested on all the items in the experiment to ensure retention. Seventeen out of 18 participants required one round of correction, the remaining participant required a second round. Participants practiced a mock version of the within scanner task on a laptop prior to scanning.

### Paradigm

In the scanner, participants were required to attend to the signed and spoken stimuli and to press a button when they encountered an item from outside the categories of fruit, animals or transport, e.g. a target filler item (see Fig. 1b). The handedness of the button press was counterbalanced across participants.

Data were collected in 6 runs. In each run, each of the 9 core items were presented twice in each of the following formats: sign and speech; male and female model. Therefore each core item was presented 8 times in each run (2×2×2), with 72 core trials in total (9 items × 8 instances). Within each run, core items were presented as two concatenated mini blocks of 36 trials. Within each mini block items were randomised with the constraint that the same concept (e.g., ‘orange’) could not be presented consecutively, regardless of modality, to reduce habituation.

In addition, in each run there were 6 target filler trials (non fruits, transport or animals) for which participants were required to press a button and 6 non-target fillers (‘other’ fruits, transport or animal items). The total number of trials was balanced within run for modality (e.g. whether sign or speech) and language model (e.g. speaker and signer). The filler trials (target and non-target fillers) were interspersed within each run regularly but unpredictably. An additional, seven null trials lasting 4 seconds were regularly but unpredictably interspersed within the each run. During these trials a white fixation cross was presented on a grey background in the absence of sound or additional visual stimulation for 4 seconds.

In summary, each of 6 runs consisted of 91 trials (72 core trials, 6 target filler trials, 6 non-target filler trials, 7 null trials). The order of modality of presentation of the items (speech/sign) was counter balanced across pairs of participants, such that items presented as signs to participant 1 were presented as speech to participant 2, and vice versa. Each stimulus was presented for its natural duration and was followed by a fixation cross lasting 3 seconds, before the start of the next trial.

After scanning, participants provided iconicity ratings on the sign stimuli that they had viewed in the scanner using the technique described by Vinson et al.^55^. They then took part in a multiple arrangement task in which they arranged pictures of the core and filler items “based on their similarity” using a drag and drop interface^58^. The Euclidean distances derived from this arrangement correlated highly with the CSLB concept property norms for the core items (r = 0.904, n = 36, p = 4.42 × 10^−14^), suggesting that the semantic feature norms provided a good summary of the semantic space of our participant group.

### Data Acquisition

Data was acquired with a 3-Tesla scanner using a Magnetom TIM Trio systems (Siemens Healthcare, Erlangen, Germany) with a 32 channel headcoil. A 2D epi sequence was used comprising forty 3mm thick slices using a continuous ascending sequence (TR=2800ms, TA=2800ms, FA= 90°, TE=30ms, matrix size= 64×64, in-plane resolution: 3mm × 3mm, interslice gap = 1mm). Six runs of data were acquired each lasting ~6-7 minutes with around 136 brain volumes collected per run; the exact number of volumes was dependent on the stimuli included in each run. EPI data collection lasted around 45 minutes. This was followed by a fieldmap, acquired using a double-echo FLASH gradient echo sixty-four slice sequence (TE1=10ms, TE2=12.46ms, in-plane view 192×192 mm, in-plane resolution: 3mm × 3mm, interslice gap = 1mm). At the end of the session a high-resolution T1 weighted structural image was collected using a 3D Modified Driven Equilibrium Fourier Transform (MDEFT) sequence (TR=1393ms, TE=2.48ms, FA= 16°, 176 slices, voxel size = 1 × 1 × 1 mm).

In the scanner, stimuli were presented using the COGENT toolbox (http://www.vislab.ucl.ac.uk/cogent.php) running in MATLAB. Auditory stimuli were presented at the same comfortable listening level for all participants. Visual images were presented using a JVC DLA-SX21 projector, with a screen resolution of 1024×768 and frame rate of 60Hz, using back projection onto a within bore screen at a distance of 62cm from the participants’ eyes.

### Univariate Analysis

Data were analysed using SPM12 (http://www.fil.ion.ucl.ac.uk/spm/). The first six images of each run were removed to account for T1 equilibrium effects. The structural and functional images were centred at the anterior commissure. Functional scans were slice time corrected to the middle slice, realigned to the first image and unwarped using field maps. The structural image was co-registered to the mean functional image. The parameters derived from segmentation, using the revised SPM12 segmentation routines, were applied to normalise the functional images that were re-sampled to 2×2×2mm. The normalized images were then smoothed with a Gaussian kernel of 6-mm full-width half maximum. Data were analyzed using a general linear model with a 360 second high-pass filter and AR1 correction for auto-correlation. In the first level design matrices, events were modelled with a canonical hemodynamic response function marking the onset of the stimulus and duration in seconds. The design matrices included a regressor for the onset of the speech trials, sign trials, filler target and non-target trials in each modality (4 regressors), button presses when the target was present in each modality (e.g. hits) (2 regressors) and button presses when the target trials were absent for each modality (e.g. false alarms) (2 regressors), six movement regressors of no interest and the session means. The rest condition constituted an implicit baseline. Contrast images of [speech > rest] and [sign > rest] were taken to the second level to conduct one sample t-tests.

### Representational similarity analysis (RSA)

At the first level, data were analysed with SPM12. Analyses were conducted in native space. Images were slice time corrected to the middle slice, realigned to the first image and unwarped using fieldmaps, but were not normalised or smoothed. The images were segmented, using the revised SPM12 segmentation routine, to estimate the transformation from native space to MNI space and vice versa. In the first level model in native space, the two repetitions of each core item presented in each modality and by each speaker and signer were modelled as a separate regressor (36 regressors: 9 core items × 2 modalities × 2 language models). Additional regressors were included modelling the onset of filler target and filler non-target trials for each modality (4 regressors), plus button presses when the target was present in each modality (e.g. hits) (2 regressors) and button presses when the target trials were absent for each modality (e.g. false alarms) (2 regressors). This constituted 42 regressors per run, plus 6 motion parameter regressors and 6 session means. A high pass filter set at 360 seconds and AR(1) correction was applied. RSA analysis was conducted with the latest version of the RSA toolbox (https://github.com/rsagroup/rsatoolbox)^59^. The representational distances estimated from the first level betas were used to calculate the cross-validated Mahalanobis (crossnobis) distances using the RSA toolbox^59^. These crossnobis distances employ multivariate noise normalisation that down-weight correlated noise across voxels, thereby increasing sensitivity to experimental effects^60^. The cross-validation across imaging runs ensures that the estimated distances between neural patterns are not systematically biased by run-specific noise, which allows us to test the distances directly against zero (as one would test cross-validated classification accuracy against chance). Therefore, the crossnobis distance provides a measurement on a ratio scale with an interpretable zero value that reflects an absence of distance between items.

A volumetric searchlight analysis^61^ was conducted using a spherical 8mm searchlight containing 65 voxels, consistent with the parameters used in previous studies of language processing^48^. In the searchlight analysis, the crossnobis distance between each core stimulus and every other was calculated to generate a Representational Dissimilarity Matrix (RDM) for every voxel and its surrounding neighbourhood. The resulting RDM reflected sign-sign, speech-speech or speech-sign distances, that constitute within and across-modality dissimilarities. In the searchlight analyses, the average of speech-speech and sign-sign distances (e.g. combined within-modality distances) and the average of the speech-speech and sign-sign distances separately were returned to the voxel at the centre of each sphere in three separate searchlight analyses. Within-modality distances were calculated only between items from the different language models (e.g. different speakers and signers respectively) to exclude similarities driven by low-level perceptual properties. Each participants’ native space whole brain searchlight map was normalised to MNI space. These maps were inclusively masked with a >20% probability grey matter mask, using the canonical MNI brain packaged with SPM12. The resulting normalised, masked images were submitted to SPM12 for one sample t-tests testing for greater than zero within-modality distances and paired t-tests testing for differences between the speech-speech and sign-sign distances at the second level. All statistical maps are presented at an uncorrected peak level threshold of p < 0.005, FDR cluster corrected at q < 0.05 to identify regions of interest for subsequent analysis.

The clusters identified from these analyses were used as Regions of Interest (ROIs) in which to test theoretical models of brain function. Note that ROI analyses are advised when testing special populations in which sample sizes are necessarily restricted^62^. Using ROIs that contain reliable representational structure, e.g. greater than zero distances, provides an additional protection against spurious distance-model correlations in regions in which there is no reliable representational structure. This approach is agnostic to the type of representational structure identified by the searchlights ensuring that ROI selection and model validation are independent from one another, and hence this does not represent “double dipping”^63^.

As each cluster contains multiple RDMs, one for each searchlight contained within the cluster, the RDMs were averaged, to provide a single representative RDM for each cluster, and each participant. These distances were then used to test hypothetical models of brain function (described below). The non-parametric Tau-a correlation was used in preference to Pearson or Spearman correlation as the models contained tied ranks^59^. The resulting correlation coefficient was converted to a Pearson’s r value, then to a Fisher-transformed Z value, to permit parametric statistical analysis^64^. Noise ceilings^59^ were estimated within-modality and across-modality separately as appropriate for each model. The lower bound was estimated by calculating the mean z converted Tau-a correlation coefficient between each participant’s RDM and the average RDM for the group excluding that participant (e.g. leaving one participant out). This is an estimate of the fit that should be achieved if the theoretical model captures all systematic variation in the RDM across subjects in this region. The upper bound was estimated by calculating the mean z converted, Tau-a correlation between each participant’s RDM and the average RDM for the group including that participant. This value constitutes a theoretical maximum of the best possible fit that can be achieved between the data and a model with this region. These limits provide a benchmark against which to assess the quality of model fit as they reflect the bounds of the best possible model fit that could be expected given the noise in the data.

### Models

A semantic model was tested using the CSLB concept property norms^28^ (Fig. 1c). This kind of feature-based semantic model can account for the ability to categorize by semantic group, e.g. a zebra is an animal, and to tell-apart unique items, e.g. that a zebra differs from a horse. As such, the similarities expressed by the model can be decomposed into two independent components. One, an **item-based** model that predicts that each item is uniquely represented, e.g., an ‘orange’ is more dissimilar to all other items than to itself, and does not predict any other relatedness between items (Fig. 1d). The other, a model in which item-to-item similarities are not tested, but category structure is predicted (Fig. 1e) – referred to as a **category-based** model. An additional model testing for dissimilarities based on speaker (Fig. 3e) and signer identity (Fig. 4e) was also tested, e.g. models predicting trials from speaker/signer 1 to be more dissimilar than trials from speaker/signer 2, and vice versa. The purpose of this model was to test for neural dissimilarities based on lower level acoustic and visual features.

These models can be tested ***within-modality***, e.g. correlated within speech-speech and sign-sign distances combined or separately, or ***across-modality***, e.g. correlated with speech-sign distances. The testing of models using ***across-modality*** distances is equivalent to cross decoding representational structure between speech and sign, positive evidence provides support for common representational structure across languages^65^. Note that we only test for ***across-modality*** semantic representations in areas in which there is evidence of ***within-modality*** representational structure. As negative correlations are not plausible, greater than 0 model fits were assessed with one-tailed, one sample t-tests. Two-tailed paired t-tests were used to assess differences in fit between models. Multidimensional Scaling (MDS) was conducted to visualise the similarity structure of the RDMs by calculating the averaged participant RDM and applying non-metric MDS, consistent with the non-parametric correlational approach.

**Table 1:** MNI coordinates for RSA analyses – 3 local maxima more than 8 mm apart

| Region | X | Y | Z | Extent | Z Value |
| --- | --- | --- | --- | --- | --- |
| **Within-modality representational structure** |  |  |  |  |  |
| Right superior temporal gyrus | 58 | -4 | -2 | 1545 | 5.283 |
| Right inferior parietal lobule | 64 | -30 | 14 |  | 4.968 |
| Right superior temporal gyrus | 52 | -2 | -8 |  | 4.861 |
| Left superior occipital gyrus | -14 | -96 | 10 | 2629 | 4.677 |
| Right superior occipital gyrus | 14 | -100 | 16 |  | 4.479 |
| Right cuneus | 6 | -92 | 22 |  | 4.226 |
| Left superior temporal gyrus | -60 | -10 | -2 | 1276 | 4.500 |
| Left middle temporal gyrus | -64 | -30 | 6 |  | 4.476 |
| Left middle temporal gyrus | -64 | -44 | 2 |  | 4.175 |
| Left inferior temporal gyrus | -48 | -62 | -6 | 172 | 4.361 |
| Left middle occipital gyrus | -42 | -64 | 0 |  | 3.122 |
| Right insula | 36 | -12 | 14 | 194 | 4.178 |
| Right putamen | 30 | -8 | 10 |  | 4.160 |
| Right middle temporal gyrus | 52 | -68 | 6 | 279 | 3.954 |
| Right middle temporal gyrus | 56 | -48 | 0 |  | 3.748 |
| Right middle temporal gyrus | 54 | -54 | 6 |  | 3.574 |
| **Greater representational structure for speech compared to sign** |  |  |  |  |  |
| Right superior temporal gyrus | 58 | -4 | -2 | 754 | 4.877 |
| Right superior temporal gyrus | 52 | 0 | -8 |  | 4.779 |
| Right superior temporal gyrus | 60 | -12 | 4 |  | 3.590 |
| Left superior temporal gyrus | -56 | -8 | 2 | 743 | 4.484 |
| Left superior temporal gyrus | -62 | -30 | 10 |  | 4.253 |
| Left superior temporal gyrus | -62 | -2 | 0 |  | 3.720 |
| Right Putamen | 30 | -10 | 10 | 146 | 4.364 |
| Right Insular | 40 | -12 | 10 |  | 3.354 |
| Right superior temporal gyrus | 58 | -34 | 18 | 285 | 4.160 |
| Right superior temporal gyrus | 66 | -32 | 14 |  | 3.763 |
| Right superior temporal gyrus | 56 | -26 | 0 |  | 3.722 |
| **Greater representational structure for sign compared to speech** |  |  |  |  |  |
| Left cuneus | -6 | -98 | 16 | 1145 | 4.623 |
| Left middle occipital gyrus | -12 | -102 | 4 |  | 4.019 |
| Left cuneus | -8 | -94 | 28 |  | 3.830 |
| Right superior occipital gyrus | 22 | -90 | 16 | 969 | 4.375 |
| Right lingual gyrus | 16 | -84 | -4 |  | 3.976 |
| Right cuneus | 16 | -100 | 12 |  | 3.655 |
| Left inferior occipital gyrus | -44 | -80 | -6 | 264 | 4.107 |
| Left middle occipital gyrus | -50 | -72 | -2 |  | 3.937 |
| Left middle occipital gyrus | -42 | -80 | 4 |  | 3.449 |
| Left cerebellum | -4 | -48 | -8 | 116 | 3.808 |
| Left lingual gyrus | -10 | -56 | -2 |  | 3.767 |
| Left cerebellum | -4 | -50 | 0 |  | 3.102 |
| Left superior occipital gyrus | -10 | -84 | 42 | 127 | 3.781 |
| Left superior occipital gyrus | -16 | -78 | 40 |  | 3.396 |
| Left superior parietal lobule | -26 | -80 | 48 |  | 3.172 |

## DATA AVAILABILITY

At the time of data collection participants did not consent to sharing their data via an open repository. Therefore, the data of this study are not publicly available. However, the data are available from the corresponding author upon request.

## ACKNOWLEDGEMENTS

This research was funded by a Wellcome Trust Senior Research Fellowship awarded to MM [100229/Z/12/Z]. CP is supported by a Wellcome Trust Principal Research Fellowship [097720/Z/11/Z]. We would also like to acknowledge support from an Economic and Social Research Council Research Centre Grant (Deafness Cognition and Language Research Centre (DCAL) [RES-620-28-0002] and a Wellcome Trust Centre Grant (203147/Z/16/Z). Thank you to Monika Grigorova and Will Dawson for help in collecting this data.

## AUTHOR CONTRIBUTIONS

S.E., M.M., J.D., C.P. & E.G. designed the study. S.E. collected the data. S.E., J.D., M.M. analysed the data. All authors contributed to writing the article.

