## Supplementary material for "Evidence for shared conceptual representations for sign and speech": All supplementary materials

**Supplementary Information 1**

During the within scanner task, on average 97% of outside category target items were identified during the semantic monitoring task (mean 35/36 correct, SD = 1.45, min = 31, max = 36) and accuracy was significantly greater than chance (mean d’ score = 4.56), t (16) = 42.74, p = 6.37 x 10^-18^.

**Supplementary Information 2**

*Contextualising the RSA findings within previous literature*

We conducted the conjunction null^66^ of [Speech > Rest] and [Sign > Rest]. As expected, areas of shared univariate activity for sign and speech were found in the bilateral posterior superior and the middle temporal gyrus, the left inferior frontal gyrus and bilateral cerebellum, consistent with previous studies^21–26,32^.

**Supplementary Fig. 1:** Areas responding to speech (red) and sign (blue) compared to rest and their overlap (pink), thresholded at p < 0.005 peak level, q < 0.05 FDR corrected at the cluster level. Rendered with MRICRON on the Ch2better brain.

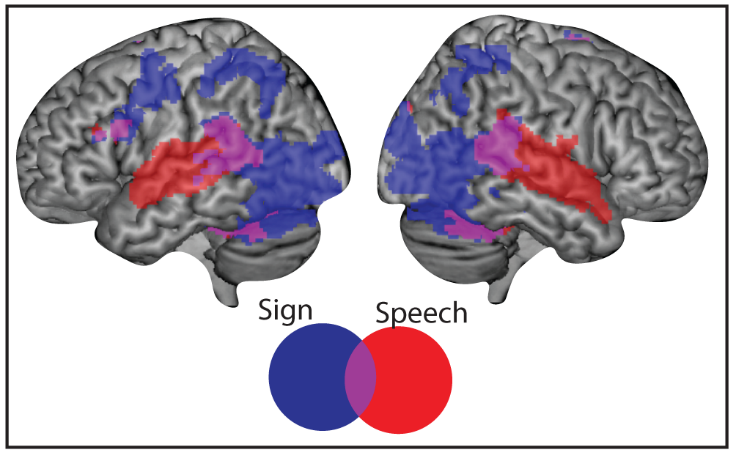

**Supplementary Information 3**

We tested the fit between the combined ***within-modality*** distances (e.g. the between speaker, combined speech-speech and sign-sign distances) and the semantic feature model (Fig. 1c) in the six clusters identified as containing reliable within-modality representational distances. We adjusted the critical alpha level to p < 0.008 to account for the six tests/clusters. Only three clusters showed a significant fit to the semantic model ***within***-***modality*** (Fig. 2b – red boxes). These were found in the right middle temporal and V5/MT (cluster 4, t (16) = 3.946, p = 5.78 x 10^-4^, d_z_ = 0.957), the bilateral V1-V3 and LOC (cluster 1, t (16) = 3.837, p = 7.28 x 10^-4^, d_z_ = 0.931) and the left posterior middle and inferior temporal gyrus (left pMTG/ITG) (cluster 6, t (16) = 3.622, p = 0.001, d_z_ = 0.879, see Fig. 2d).

*Within-modality effects for sign not speech*

We further tested for a difference in the strength of fit of the semantic feature model to the sign compared to speech distances, in the three clusters that showed a significant fit to the within modality semantic feature model. We adjusted the critical alpha level to p < 0.017 to account for the three tests for the following tests.

The clusters within the visual cortices showed a stronger fit to the semantic model for sign than for speech: the right middle temporal and V5/MT cluster (t (16) = 2.842, p = 0.012, d_z_ = 0.689) and the bilateral V1-V3/LOC cluster (t (16) = 4.630, p = 2.78 x 10^-4^, d_z_ = 1.123). Follow up tests in both regions, showed a significant fit to the semantic feature model for sign (both ps < 1.05 x 10^-4^) but not for speech (both ps > 0.110).

*Within-modality effects for both sign and speech*

By contrast, the cluster in the left pMTG/ITG did not show evidence of a difference in the strength of semantic encoding between speech and sign in the ***within***-***modality*** distances (t (16) = 0.400, p = 0.694, d_z_ = 0.097). The semantic feature model was a marginally significant fit to both the speech-speech (t (16) = 2.188, p = 0.022, d_z_ = 0.531) and sign-sign (t (16) = 2.195, p = 0.022, d_z_ = 0.532) distances separately.

*Across-modality effects for both speech and sign*

Finally, we tested for a fit to the semantic feature model in the across-modality distances. Only the response in the left pMTG/ITG was a significant fit to this model (t (16) = 3.076, p = 0.004, d_z_ = 0.746), at an adjusted alpha of p < 0.017 (see Fig. 2d).

**Supplementary Information 4**

We used the participants’ average iconicity rating for each sign to generate an iconicity model (Supplementary Fig. 2, below) to test for an influence of iconicity on neural responses. This model was created by taking the absolute value from the subtraction of the average iconicity value of each sign from the iconicity value of every other sign. We tested for a fit between this model and the neural response in the pMTG/ITG. This showed that there was no significant fit to the within sign (t (16) = 0.382, p = 0.354, d_z_ = 0.092) or **across-modality** distances (t (16) = 1.298, p = 0.106, d_z_ = 0.315).

**Supplementary Fig. 2:** Iconicity model derived from participant ratings of the iconicity of the sign stimuli (left). The model (right) was tested on the sign-sign distances (red box), e.g. within sign, and the speech-sign distances (blue box), e.g. across-modality.

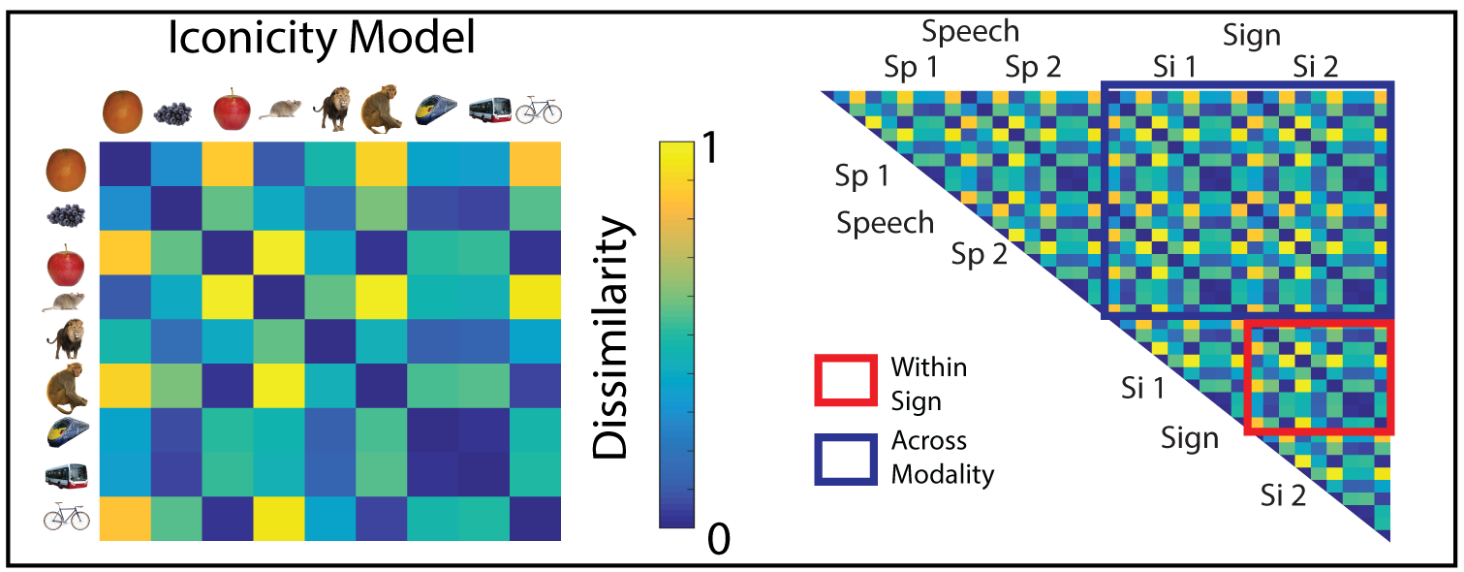

**Supplementary Fig. 3**: tSNR maps showing sagittal slices of the left temporal lobe.

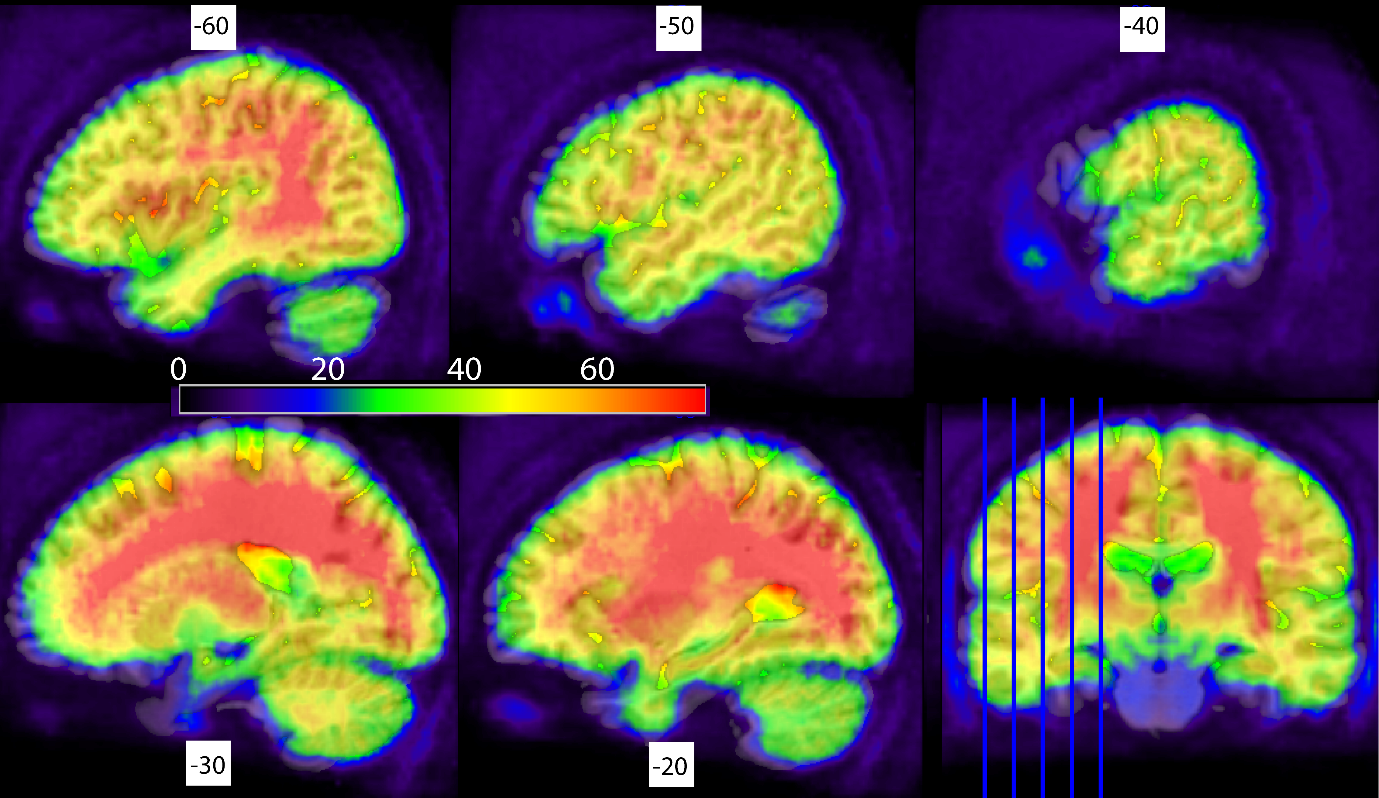
